# Persistent horizontal and vertical, MR-induced nystagmus in resting state Human Connectome Project data

**DOI:** 10.1101/2022.02.23.481619

**Authors:** Cammille C Go, Huseyin O Taskin, Seyed-Ahmad Ahmadi, Giulia Frazzetta, Laura Cutler, Saguna Malhotra, Jessica IW Morgan, Virginia L Flanagin, Geoffrey K Aguirre

## Abstract

**Objective:** Strong magnetic fields from magnetic resonance (MR) scanners induce a Lorentz force that contributes to vertigo and persistent nystagmus. Prior studies have reported a predominantly horizontal direction for healthy subjects in a 7 Tesla (T) MR scanner, with slow phase velocity (SPV) dependent on head orientation. Less is known about vestibular signal behavior for subjects in a weaker, 3T magnetic field, the standard strength used in the Human Connectome Project (HCP). The purpose of this study is to characterize the form and magnitude of nystagmus induced at 3T.

**Methods:** Forty-two subjects were studied after being introduced head-first, supine into a Siemens Prisma 3T scanner. Eye movements were recorded in four separate acquisitions over 20 minutes. A biometric eye model was fit to the recordings to derive rotational eye position and then SPV. An anatomical template of the semi-circular canals was fit to the T2 anatomical image from each subject, and used to derive the angle of the B_0_ magnetic field with respect to the vestibular apparatus.

**Results:** Recordings from 37 subjects yielded valid measures of eye movements. The population-mean SPV ± SD for the horizontal component was −1.38 ± 1.27 deg/sec, and vertical component was −0.93 ± 1.44 deg/sec, corresponding to drift movement in the rightward and downward direction. Although there was substantial inter-subject variability, persistent nystagmus was present in half of subjects with no significant adaptation over the 20 minute scanning period. The amplitude of vertical drift was correlated with the roll angle of the vestibular system, with a non-zero vertical SPV present at a 0 degree roll.

**Interpretation:** Non-habituating vestibular signals of varying amplitude are present in resting state data collected at 3T.

## Introduction

Strong static magnetic fields, such as those produced by a magnetic resonance (MR) imaging scanner, can induce vestibular signals that may be observed as nystagmus. This phenomenon accounts for the sensation of rotation and vertigo that people frequently experience within MR scanners (Friebe et al., 2015; Glover et al., 2007; Mian et al., 2013; Ward et al., 2014). The availability of visual input allows people to suppress this nystagmus, despite persistence of the vestibular signal. Roberts and colleagues proposed that MR-associated vertigo and persistent nystagmus occur due to a Lorentz force produced by the interaction of static magnetic fields and ionic currents in inner ear endolymph fluid (Roberts et al., 2011). This pressure stimulates the semicircular canal cupula and contributes to a sensation of head rotation, with the resultant force scaling linearly with the strength of the magnetic field (Antunes et al., 2012; Ward et al., 2018).

Prior studies have characterized the form and magnitude of slow-phase velocity (SPV) eye movements within a 7T MRI scanner, with a specific focus on the relationship between head position and the direction and magnitude of nystagmus. The predominant effect reported has been horizontal SPV that is dependent upon degree of head pitch (nose up, nose down) (Boegle et al., 2016; Roberts et al., 2011). Vertical nystagmus has been related to head roll (tilt left, tilt right), although the magnitude of the vertical as compared to horizontal SPV has been reported to be relatively small (Mian et al., 2016). As these data were primarily collected using 7T MRI scanners, there has been less examination of distribution of SPV (and thus vestibular signals) present within a typical, healthy population of people studied in 3T scanners (Boegle et al., 2016). The Human Connectome Project (HCP) has collected measurements from several thousand people under “resting state” conditions at 3T (Glasser et al., 2016). To better characterize resting-state data, it would be useful to know the form and individual distribution of vestibular signals that are present within these measurements.

In this study, we measured MR-induced, SPV eye movements in people who were in complete darkness. The measurements were made on a 3T scanner, using typical HCP hardware and protocols. The magnitude of horizontal and vertical SPV eye movements was measured in four separate acquisitions over a 30-minute period. We examined the distribution of nystagmus within this population, exploring the influence of head position upon our results. Our findings indicate that there are substantial, and varying, horizontal and vertical vestibular signals that persist throughout the recording period.

## Materials and Methods

### Participants

We studied 42 participants from the University of Pennsylvania and the surrounding Philadelphia community as part of a pre-registered project (https://osf.io/ervrh). These subjects served as normally-sighted controls for the HCP “Human Connectomes in Low Vision” project. Participants were at least 18 years of age, had no history of ophthalmologic disease, and had corrected visual acuity of 20/40 or better. Fourteen of the subjects were women, and the mean age of all participants was 28. This study was approved by the University of Pennsylvania Institutional Review Board, and all participants provided written informed consent. This work was carried out in accordance with the Declaration of Helsinki.

### Scanning environment and MR imaging of the brain

At least two MR imaging sessions were conducted for each subject. The data presented here are drawn from the first session, which focused upon structural imaging and “rest state” functional MRI. During this first session, all sources of light within the scanner room were extinguished and covered. The complete absence of light within the room was confirmed by having some of the investigators dark adapt for 30 minutes, and then systematically pursue and cover any source of light. Room lights were on when subjects were positioned within the scanner, but these were turned off at the start of data acquisition. Room lights were turned on later during the session for the collection of structural images and during “gaze calibration” eye tracking (described below).

As part of our scanning protocol, we asked subjects at the half-hour point of their session if they were able to perceive any sources of light. Three of the 42 subjects reported being able to dimly perceive the active, infra-red LED of the eye tracking camera. They described this sensation as colorless, and in the periphery of their vision, consistent with reports of rod-mediated perception of light with wavelengths longer than 800 nm (Griffin et al., 1947). In our analyses we considered the possibility that these subjects had a source of visual fixation that might attenuate their nystagmus.

MRI scans made use of the HCP LifeSpan protocol (VD13D) implemented on a 3 Tesla Siemens Prisma with a 64-channel Siemens head coil. Subjects were placed supine on the scanner table, and introduced into the scanner head-first. BOLD fMRI data were obtained over 72 axial slices with 2 mm isotropic voxels with multi-band = 8, TR = 800 ms, TE = 37 ms, FOV = 208 mm, flip angle = 52°. We also collected a T2 image with 0.8 mm isotropic voxels, repetition time (TR) = 3200 ms, echo time (TE) = 563 ms, flip angle = 120°, FOV = 240 x 256. We attempted to collect four, resting-state BOLD fMRI acquisitions in each subject. Resting state data collection began 15 – 20 minutes after the start of the scanning session. A TTL pulse produced by the scanner at the start of each BOLD acquisition was recorded and used to synchronize the fMRI and eye video data.

The B_0_ magnetic field of the Siemens Prisma scanner has the south pole at the end at which the subject is introduced into the scanner. As our subjects were introduced into the scanner head-first, the magnetic field vector points from the feet towards the head of the subject.

### Eye video recording

An MR-compatible, IR video camera (LiveTrack, Cambridge Research Systems) was placed behind an angled cold mirror, and both the camera and mirror were mounted to the head coil. The mirror both hid the camera and allowed the subject to view a stimulus screen that was placed at the end of the magnet bore. Subjects viewed stimuli presented on an MR-compatible LCD screen (SensaVue fMRI, Invivo corporation) at an optical distance of ~1065 mm from the eye of the subject. The camera had one active, IR LED (peak emission at 940 nm), located 14 mm lateral to the center of the camera lens. Recordings of the eye were made in NTSC DV 30Hz, and in subsequent processing converted to progressive 60 Hz videos, using “bob” deinterlacing, yielding a final image size of 640 x 480 pixels. The intrinsic matrix and radial lens distortion for the camera were measured using the MATLAB camera calibrator application (Bouguet, 2012).

Two types of eye recordings were obtained. During “gaze calibrations”, the subject was asked to fixate upon targets presented on the screen. There were nine targets, arranged in a square, 3×3 grid that was 14° wide. The targets were presented in a random order and the subject was asked to fixate upon each target for several seconds. A gaze calibration measurement was obtained at the end of the first scanning session, and a further set of 1-to-4 gaze calibration measurements were made during the second scanning session.

The second type of eye recording was made during each of four “resting” fMRI scans of 336 seconds duration. Subjects were instructed only to stay awake and keep their eyes open during this measurement. Following our pre-registered protocol, scanning was stopped if the subject was observed to close their eye for longer than 5 seconds, or if there was visible movement of the head. We then attempted to re-collect the scan.

### Measurement of eye position and the velocity of slow phase drift

The IR videos of the eye were first analyzed to extract eye features using open-source software (https://github.com/gkaguirrelab/transparentTrack). Points on the border of the pupil were identified, as was the center of the reflection of the active IR light source from the tear film (the “glint”). Next, the set of gaze calibration measurements for each subject, taken from both sessions, were analyzed to derive a subject-specific biometric model of the eye (Aguirre, 2019). Finally, this biometric model was used to fit the pupil and glint features on each frame of the eye videos collected during resting-state fMRI. This last step yields a measure of the horizontal and vertical rotation of the eye in degrees. Positive values in the horizontal direction indicate a rightward drift whereas those in the vertical direction indicate upward drift.

The eye-position time-series data from each acquisition was filtered to remove those frames with a poor measurement of eye position. A frame of the video was retained if:

1. A glint was present in the image
2. The elliptical model fit of the pupil perimeter points had a root mean square error (RMSE) of less than 2.25 pixels
3. The change in estimated eye position from the prior frame was less than 100 deg / second.

These criteria were designed principally to remove frames in which the pupil was obscured by blinks. Because our chief interest in this work is the measurement of slow phase eye movements, we used a velocity threshold that risked the rejection of some true, high velocity saccadic movements in exchange for improved rejection of artifactual velocity measures at the onset and offset of blinks. For some subjects, poor contrast between the pupil and iris in the IR video also produced a high level of noise in the extraction of the pupil perimeter points, causing many video frames to be rejected.

If, for a given acquisition, more than 80% of frames were rejected by these criteria, then the entire acquisition was rejected. If a subject had fewer than three remaining acquisitions to analyze, the subject was excluded from this study. Of the 42 subjects studied, one was removed because they did not have at least 3 rest-state acquisitions in a single session, and an additional four subjects were removed due to poor quality eye tracking.

In the time-series measurements of eye position that remained, we fit a linear slope model to the horizontal and vertical eye position in each acquisition to estimate the slow phase velocity drift of the eye. Horizontal and vertical drift were assessed independently. The model fit was applied iteratively as follows. First, the filtered eye-position vector was segmented using the MATLAB *ischange* function (Killick et al., 2012). A linear fit was then made to each of the segments over time and the median slope across the segments retained. The median slope is an estimate of the SPV. We then asked how well this single SPV value would account for the data, expressed as the RMSE of the residual eye position data within the segments after removing the median slope effect. The segments with the top 2.5% RMSE were then discarded, and the analysis was repeated. A set of 10 iterations was performed, causing the analysis to converge upon a final SPV value for eye movement in the acquisition. This analysis presumes a constant SPV across the duration of an acquisition, which appears well supported given our finding that SPV does not differ across acquisitions (described in Results).

The analysis for each subject across the multiple acquisitions yielded a set of 3 or 4 SPV values each in the horizontal and vertical directions. The covariance matrix for this set of measurements was used to define a bivariate error ellipse. We made use of the 95% confidence interval (CI) for inference, and plotted the 67% CI (AJ Johnson, 2022), which corresponds to the standard error of the mean.

### Calculation of brain rotation within the scanner

The field-of-view for the T2 image (including oblique rotations) was set using the autoAlign feature of the Siemens scanner, which calculates a 3D affine registration between an initial, scout image of the subject and a template brain. The template brain is an atlas that is placed in a standard orientation, including aligning the anterior-posterior commissure line (AC-PC) with the axial plane (corresponding to Talairach orientation). The *qform* matrix in the NIFTI header of the T2 image describes the transformation between image and magnet coordinates.

For each session we derived the Euler angles (pitch, roll, and yaw) of the brain of the subject (as registered to the template brain) relative to the scanner bore, and thus the B_0_ magnetic field. A nuance of this measurement is that pitch is expressed based upon the position of the brain, as opposed to (e.g.) skull based landmarks, such as Reid’s plane. Six subjects had their structural images collected during an imaging session separate from that in which their resting state fMRI data were collected. Therefore, the data from these subjects were omitted from the subsequent analyses as the orientation calculated from these images would not necessarily reflect the orientation of the head at the time of resting state data collection.

### Calculation of semicircular canal rotation within the scanner

We also calculated the rotation of the vestibular apparatus in each subject with respect to the scanner coordinates and thus the B_0_ magnetic field. To do so, we first diffeomorphically registered an MRI atlas of the human vestibular system (Ahmadi et al., 2021) to the T2 image from each subject using ANTs (Avants et al., 2011). The atlas outlines each of the six SSCs with a set of fiducial markers defined in the magnet coordinates of the image. The registration then moves these fiducial markers to the magnet coordinates of the subject T2 image. A set of five markers were defined around the perimeter of each SCC. A plane was fitted to the five markers via least-squares estimation, yielding the plane normal. In three subjects the diffeomorphic registration produced an invalid, non-anatomical result. We were unable to correct this registration error and thus, in addition to the six subject who did not have structural images collected along with their resting state data, we excluded these subjects from subsequent analyses concerning the effect of orientation of the vestibular system.

We combined the measurements across subjects to create a model of the SCC normals. To do so, we rotated the set of normals from each subject to best align with the x-y-z axes using the Kabsch algorithm (Kabsch, 1976), and then averaged the set of normals across subjects for each of the six SCCs. This produced a population-average model of the relative angles between the normals of the six SCCs.

Finally, using the same algorithm we calculated the Euler angles that produced the best match between the average, SCC vector model and the set of vectors measured for each subject. We took the resulting values of pitch, roll, and yaw as capturing the rotation of the vestibular apparatus in each subject with respect to the scanner coordinates, and thus the B_0_ magnetic field.

## Results

### Horizontal and vertical nystagmus is present in a majority of subjects within a 3 Tesla MRI

We collected IR video of the right eye of 42 subjects in a 3T MRI scanner, with data from 37 subjects retained in the final analysis. Analysis of the video provided the horizontal and vertical rotation of the eye over time (Figure 1). The static field of an MRI scanner can stimulate the vestibular apparatus of the inner ear, inducing a slow drift of the eyes (Roberts et al., 2011). People respond to this involuntary drift by engaging in corrective saccadic eye movements, resulting in nystagmus and a characteristic saw-tooth pattern of eye position over time. We fit the eye position data from each of several, 336 second acquisitions from each subject to measure the SPV movement of the eye in the horizontal and vertical direction.

**Figure 1.**
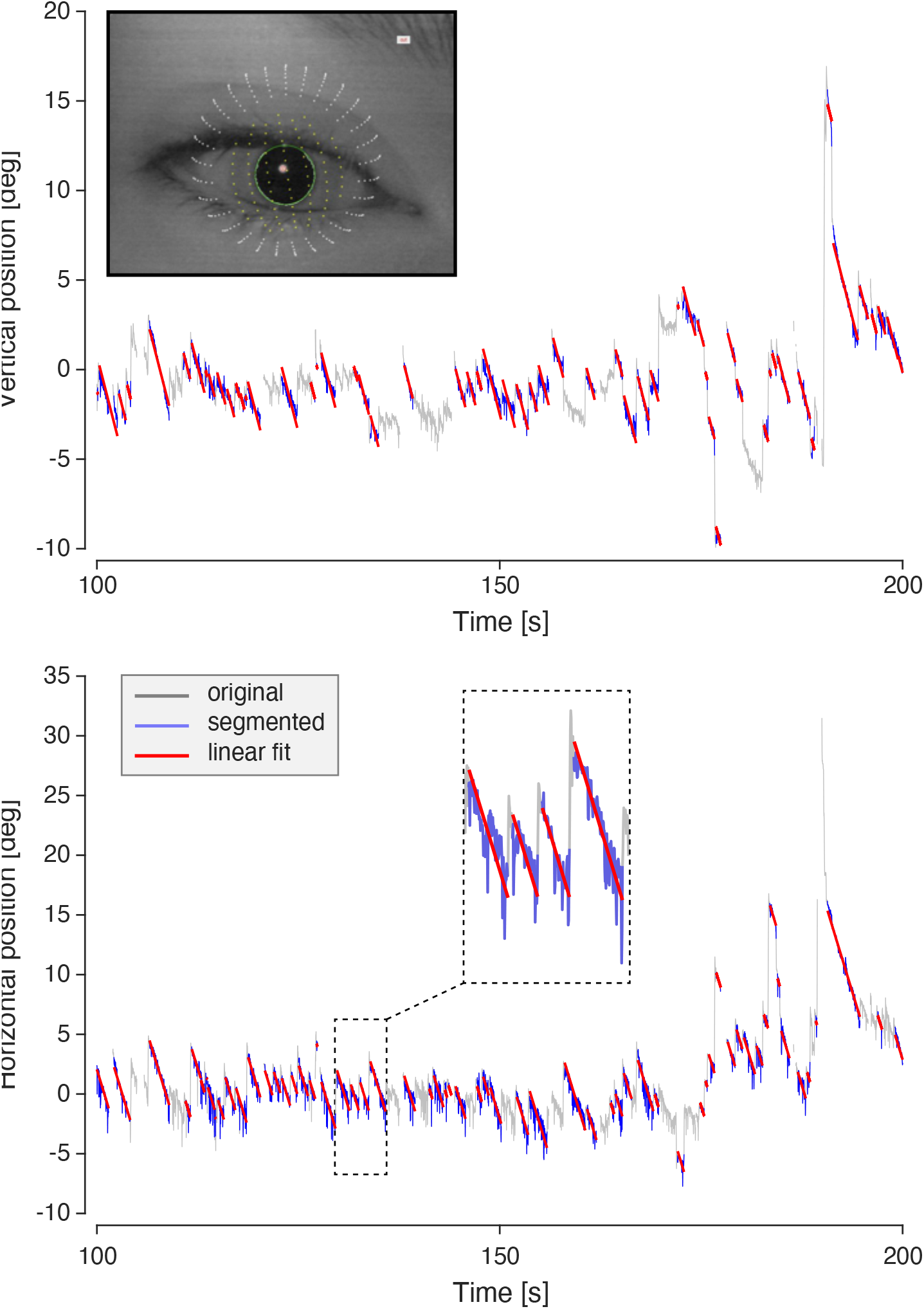
Example of linear fitting for nystagmus. An example of the horizontal (*top*) and vertical (*bottom*) nystagmus response in a single eye movement recording acquisition for a subject (TOME_3008) over 100 seconds. Shown inset (*top*) is the biometric eye model fit to a frame of video. Raw eye position data (grey) was filtered for blinks, eye-tracking artifacts, poor fitting, and outliers. After segmentation, linear fitting, and iterated removal of the segments with the highest root mean square error, the retained data (blue) was used to determine the median slope for the acquisition (red). Shown inset (*bottom*) is an expanded view of a portion of these data.

Figure 2 summarizes the SPV movement recorded from this population. Overall, subjects experienced horizontal and vertical nystagmus of similar magnitude, with a population mean (± SD) SPV of −1.38 ± 1.27 deg/sec horizontal, and –0.93 ± 1.44 deg/sec vertical. These values correspond to a slow drift of the eye rightward and downward (from the subject perspective). We calculated the confidence interval of the SPV measured in each subject over their 3 or 4 acquisitions. Nineteen subjects had a component of horizontal drift for which the 95% CI differed from zero, and the same number had a vertical drift that differed from zero. Three subjects reported being able to perceive the IR LED of the video camera, and the data from two of these subjects were retained in the final analysis. The SPV values measured for these two subjects (TOME_3013: −2.71 deg/sec horizontal, 0.30 deg/sec vertical; TOME_3046: −0.20 deg/sec horizontal, −1.63 deg/sec vertical) did not differ notably from the mean effects recorded for the population, suggesting that this minimal, peripheral perception of light was not sufficient for them to extinguish their nystagmus.

**Figure 2.**
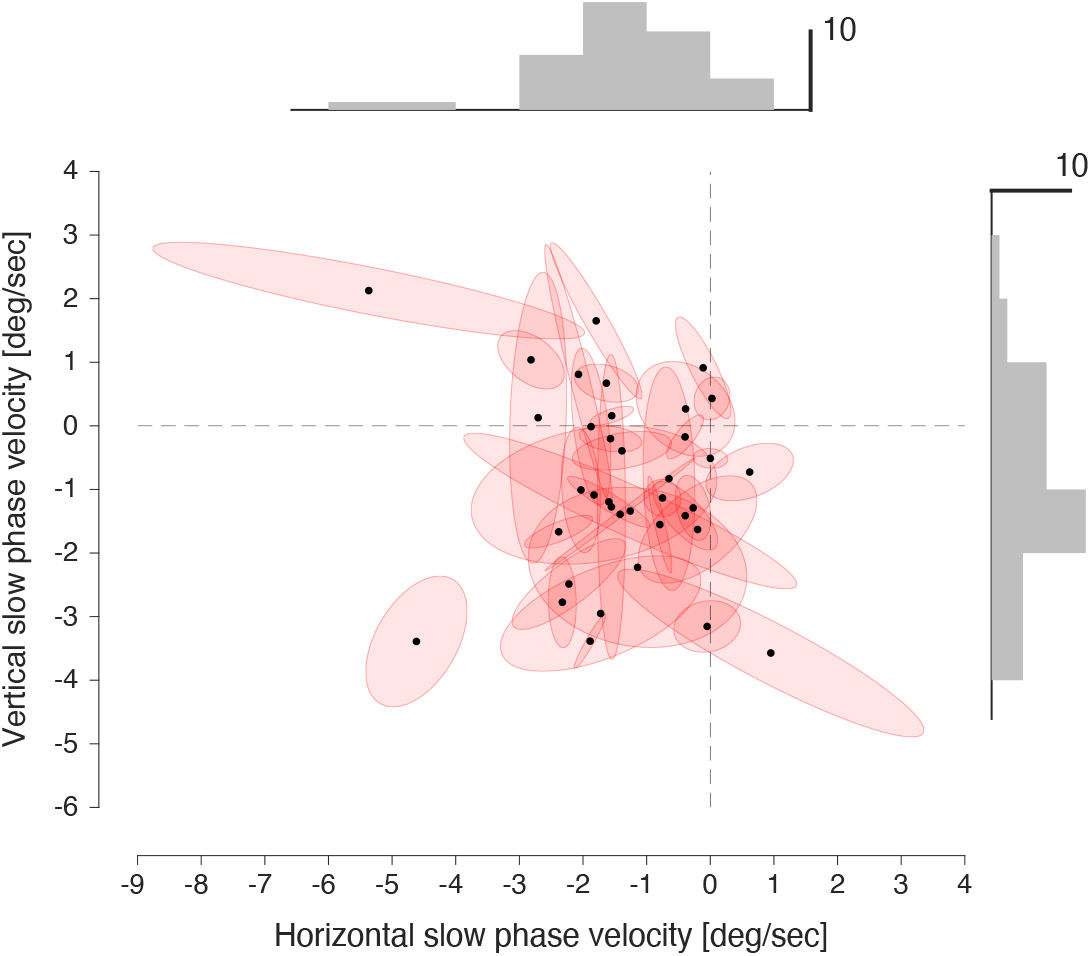
Robust horizontal and vertical slow-phase velocities. Horizontal and vertical SPV measurements in a population of 37 subjects. Each black data point is the measurement from a single subject averaged over 3 or 4 acquisitions. The bivariate ellipse around each point is the 67% confidence interval derived from the multiple acquisitions. Marginal histograms demonstrate that the central tendency of the population is to a slight rightward and downward SPV movement.

We note that the polarity of the MRI scanner used in our studies is reversed from that of several prior studies (Roberts et al., 2011; Ward et al., 2014). Correspondingly, the direction of the central tendency of the SPV effect that we observe in our data is the reverse of that reported in these prior measurements.

### A measure of the rotation of the vestibular apparatus with respect to the magnetic field

We wished to examine the relationship between individual differences in SPV eye movements and the orientation of the semicircular canals (SCCs) with respect to the B_0_ magnetic field of the scanner. We registered an MR image atlas of the left and right vestibular apparatus to the T2 image from each subject (Figure 3a). This yielded the vectors that are normal to the three SSC for each inner ear (Figure 3b). We were able to obtain this measurement in 28 of our subjects. The angles of these vectors relative to one another were combined across subjects (Figure 3c). Consistent with prior measurements (Della Santina, 2005) the angles between the SSC normals were roughly 90 degrees, with some variability across subjects and ears due to a possible combination of individual differences and errors related to image registration (left/right ear values lateral-anterior: 88.7 ± 7.2, 86.6 ± 7.4; lateral-posterior: 89.3 ± 4.3, 89.6 ± 4.9; anterior-posterior: 89.5 ± 6.5, 87.9 ± 8.6).

**Figure 3.**
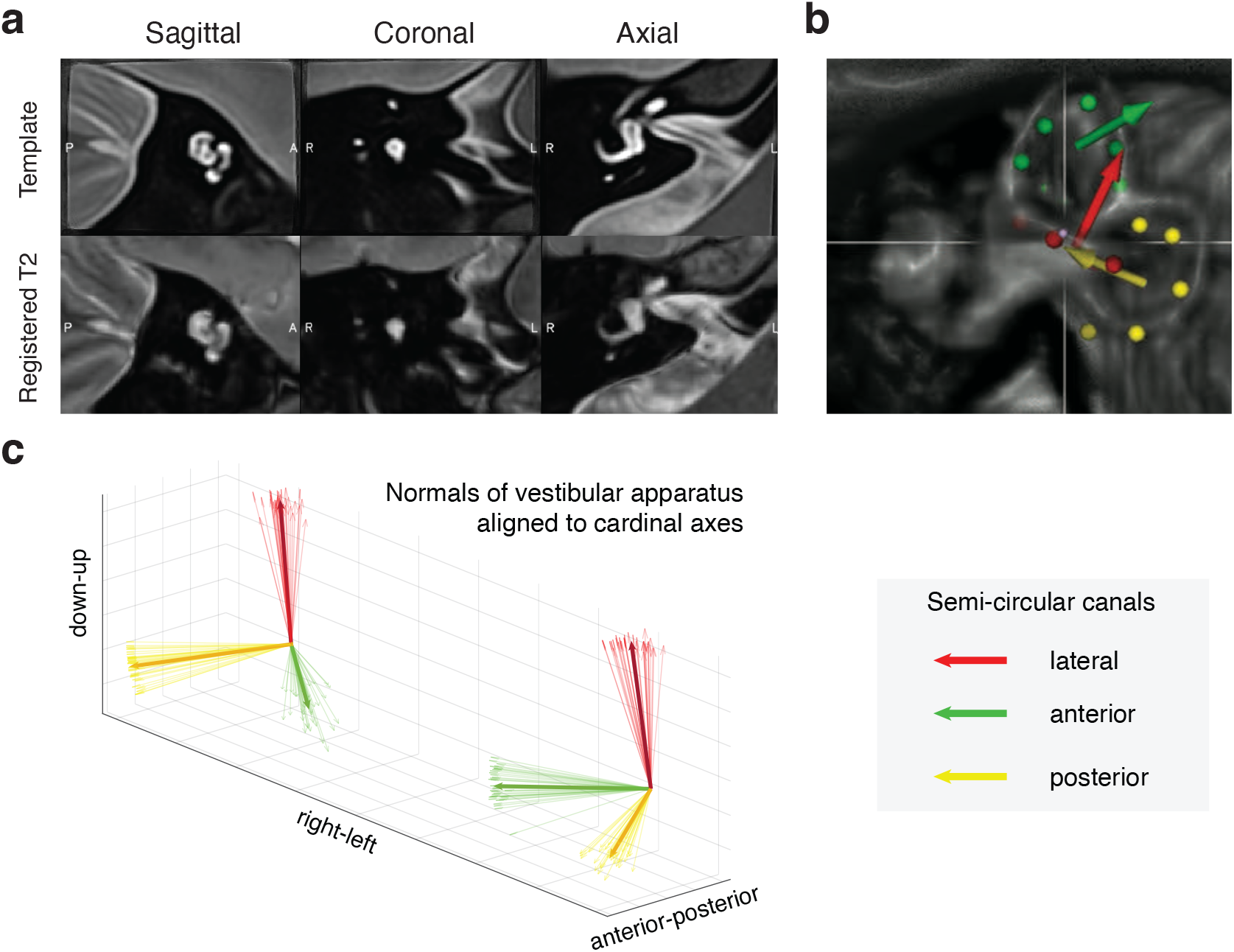
Measurement of the orientation of the vestibular apparatus. (a) *top row*: sagittal, coronal, and axial views through the left inner ear of the Ahmadi et al., 2021 atlas. The cochlea is seen, as is a portion of the lateral semi-circular canal (SCC) in the axial view. *bottom row*: corresponding views from the T2 image from one subject following registration to the atlas. (b) The atlas defines points on the plane of each SCC, and the vector that is normal to the plane. (c) The set of six SCC vectors from each subject were all aligned to the cardinal axes. Each thin arrow is a measurement from one subject. The distribution of vectors reflects variation in the relative angles between the SCCs across subjects. The mean vector across subjects is indicated by the heavier, darker line.

We calculated the Euler angles that brought this set of mean SSC normals into alignment with the set of SSC normals measured in each subject. This provides an estimate of the pitch, roll, and yaw angles of the vestibular apparatus in each subject with respect to the B_0_ magnetic field of the scanner. We also calculated for each subject the Euler angles of the brain with respect to the B_0_ field, as derived from the pulse sequence auto-align function. We first confirmed that our calculation of vestibular angles and brain angles were in general agreement (Figure 4). Across subjects, there was a high degree of agreement between the vestibular and brain angle measurements for roll and yaw, with the best fit line passing through [0,0]. This is to be expected as the mirror symmetry of the head leads to the left and right vestibular apparatus having a roughly fixed position with respect to the left and right sides of the brain. A somewhat less systematic relationship was observed between pitch as measured for the vestibular apparatus and as measured for the brain. The left and right inner ear can be rotated about the left-right axis without breaking symmetry with respect to local structures, allowing for this individual difference.

**Figure 4.**
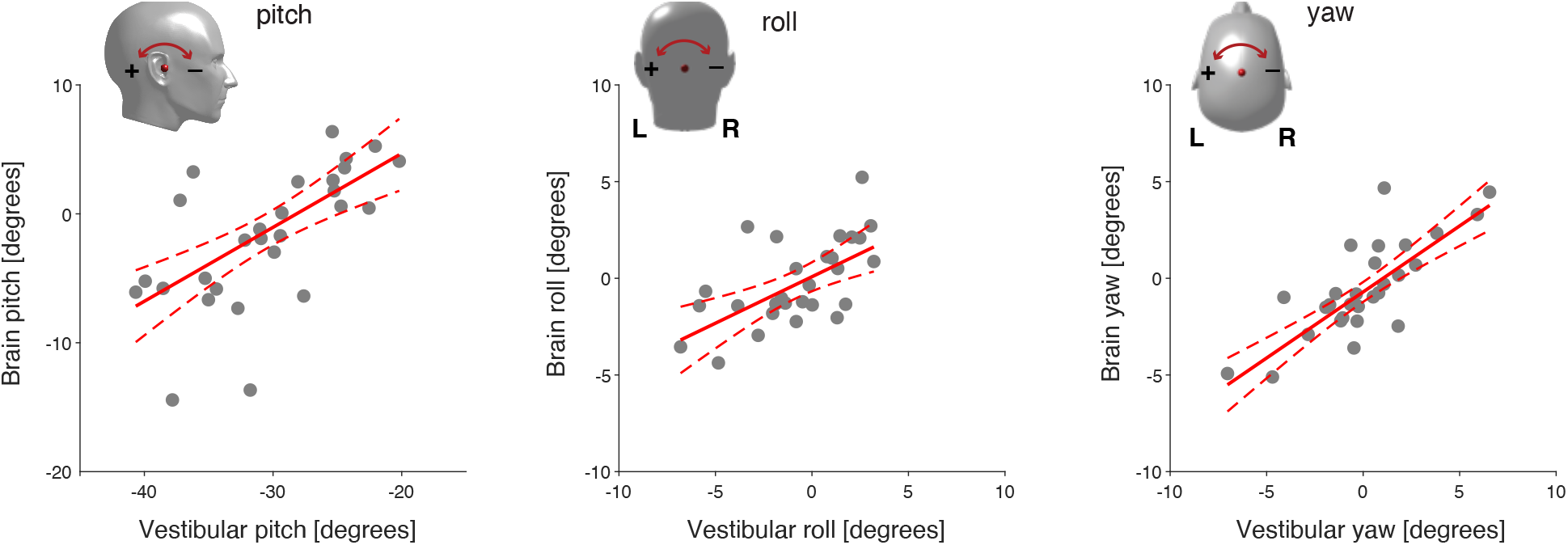
Rotation with respect to the B_0_ field calculated for the brain and vestibular apparatus. For each subject we derived the orientation of the brain, and the orientation of the vestibular apparatus, with respect to the B_0_ magnetic field. We found that the two measures are strongly related for yaw and roll, but less so for pitch. The red line is the linear fit, with the 95% confidence interval indicated by dotted lines.

We next examined if variation in these rotation angles could explain variation in the degree of SPV eye movements during scanning.

### Vertical nystagmus is related to roll angle of the vestibular apparatus

Previous work has demonstrated that head orientation with respect to the B_0_ field influences the direction and magnitude of MRI-induced nystagmus (Mian et al., 2016; Roberts et al., 2011; Ward et al., 2018). We therefore examined the across-subject relationship between the estimated rotation angles of the vestibular system and SPV eye movements.

Following prior work, we examined the relationship between pitch angle of the vestibular apparatus and horizontal SPV (Figure 5a). We did not observe a systematic relationship (slope 0.00; 95% CI −0.04 to 0.04). We examined the relationship between brain pitch angle and horizontal SPV, and again did not observe an effect (slope −0.04; 95% CI −0.09 to 0.02).

**Figure 5.**
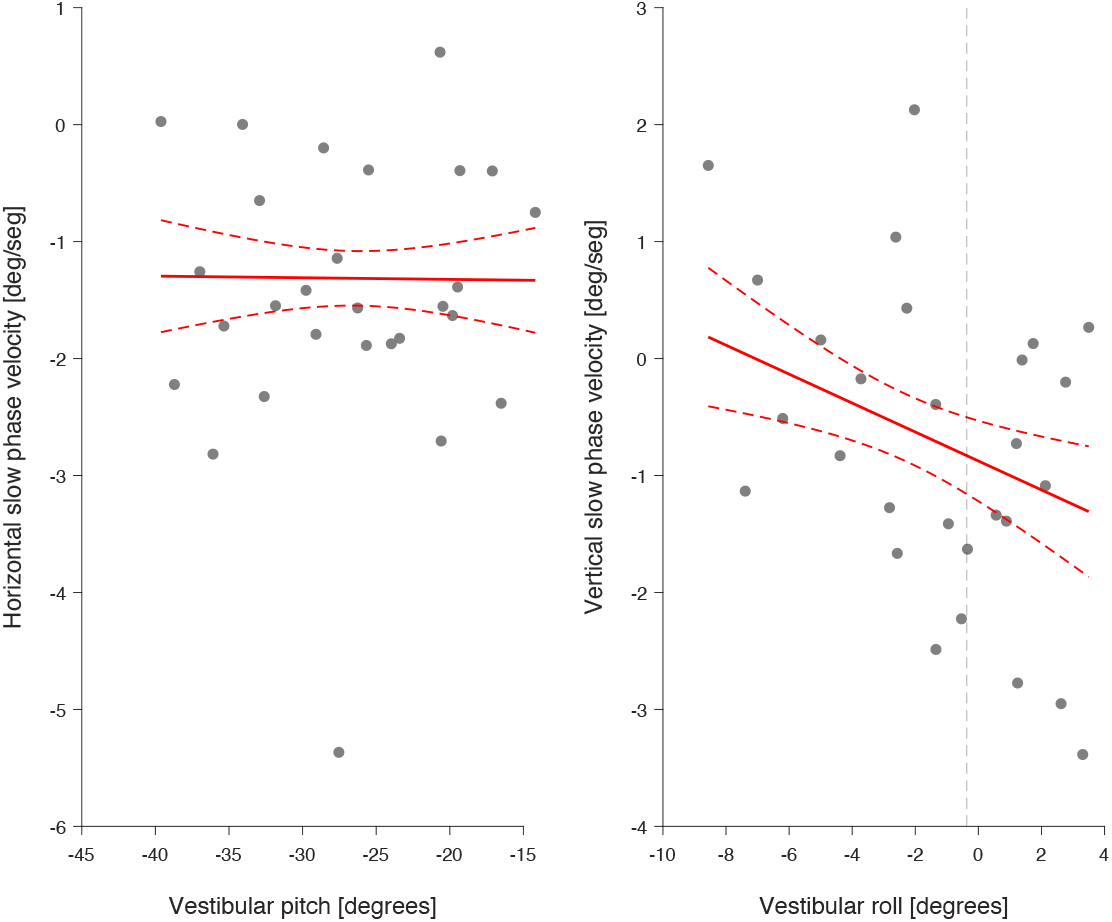
Vestibular angle with respect to B_0_ field related to slow phase velocity eye movements. Distribution of horizontal and vertical slow phase velocities (SPVs) plotted against vestibular pitch and roll, respectively. The trend of SPVs was determined through linear fitting of the mean SPVs for each subject weighted by the reciprocal of the standard deviations for each mean. Dotted red lines depict the 95% confidence interval.

In contrast, we found that the roll angle of the vestibular system was related to individual differences in vertical SPV (Figure 5b; slope −0.15; 95% CI −0.25 to −0.05). A linear fit to the relationship shows that −0.95 deg/sec of vertical nystagmus would remain (on average) even with the vestibular system positioned at a neutral position of 0 degrees roll. Similar results were obtained when these calculations were performed using the calculated brain angle instead of vestibular angle (slope −0.15, 95% CI −0.27 to −0.02).

No relationship was observed between yaw and either horizontal or vertical SPV.

### No evidence for habituation of nystagmus over 20 minutes

Three or four measurements of SPV were obtained for each subject from a recording period of over 20 minutes. We examined if there was a tendency for the amplitude of SPV eye movements to decline over the duration of the recording period. Figure 6 presents the population mean SPV value for horizontal and vertical eye rotation as a function of acquisition number. There was no tendency for SPV values to decrease over the measurement period.

**Figure 6.**
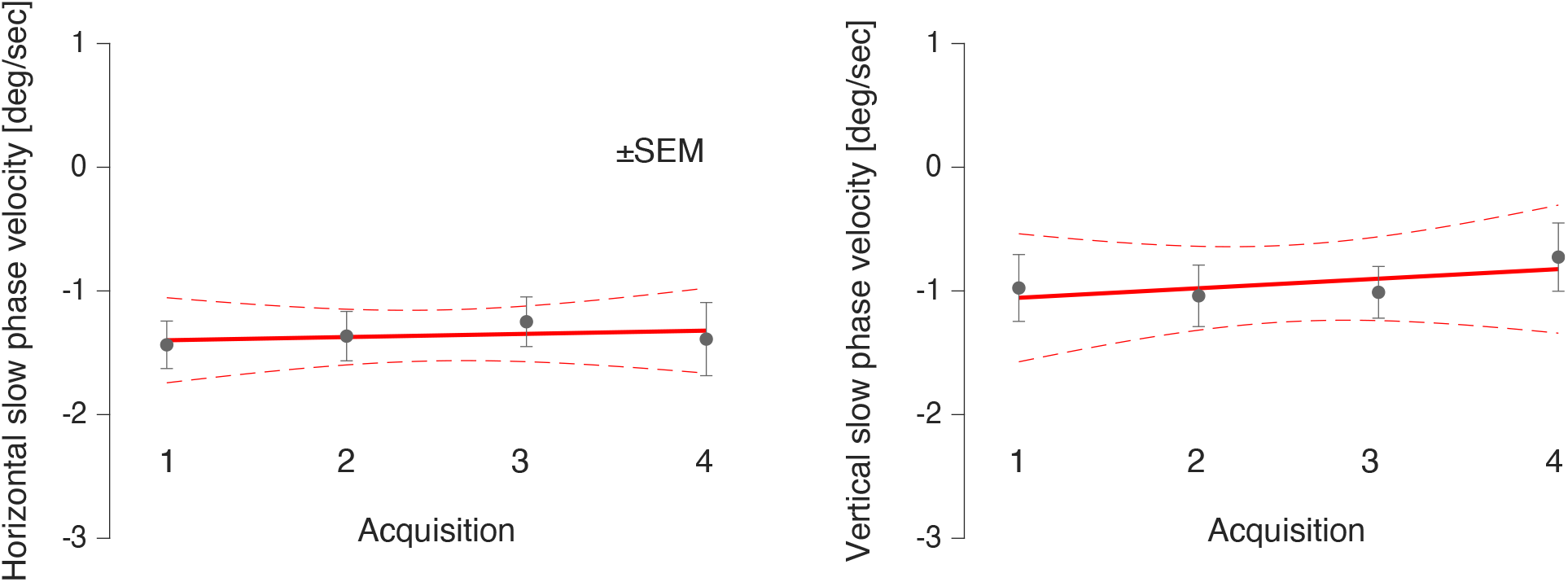
Nystagmus response persists across acquisitions. Each dot depicts the mean of slow phase velocities (SPVs) for each sequential acquisition across subjects. Error bars denote 1 standard error of the mean. Dotted red lines indicate the 95% confidence interval. The slope (95% confidence interval) for horizontal and vertical SPVs over time is 0.03 (−0.16 to 0.21, p = 0.606) and 0.08 (−0.20 to 0.35, p = 0.353), respectively.

## Discussion

We find persistent nystagmus in half of subjects studied during HCP resting-state MRI scanning at 3T. The magnitude of the effect varies substantially between subjects, and is predicted to be present (in the case of vertical nystagmus) even when the head is perfectly aligned with the B_0_ magnetic field. Given that subjects were positioned in the scanner so that they could subsequently view a stimulus screen, we expect that these vestibular signals are characteristic of all 3T HCP resting-state data, even when frank nystagmus is suppressed by visual fixation.

Prior analyses in 7T MRI scanners have generally emphasized the horizontal component of nystagmus. When horizontal and vertical SPV have been measured, the horizontal component has been observed to be larger (Glover et al., 2014; Mian et al., 2016, 2013; Otero-Millan et al., 2017). In our study we find a population-mean horizontal SPV that is smaller than previously reported, even accounting for the lower B_0_ magnetic field strength in the current measurements. A potential explanation is that prior studies positioned the head of the subject to enhance horizontal nystagmus so as to better study the effect. This is plausible as there are a variety of head pitch angles which are comfortable within a standard head coil, but a more limited range of roll angles.

We did not observe a relationship in our data between vestibular pitch and horizontal SPV across individuals. This correlation has been demonstrated previously within subject (Roberts et al., 2011), when head angle has been varied as much as +40 and −40 (Mian et al., 2016). At the extremes of head pitch, the direction of the horizontal SPV switches from left to right. A head pitch angle may be found that nulls the horizontal SPV, but this particular angle varies substantially between subjects (Roberts et al., 2011), perhaps because of individual variation in the rotational orientation of the semicircular canals in relation to external skull landmarks. The absence of a relationship between pitch and horizontal SPV in our data is likely a result of these individual differences, as well as the smaller range of head pitch angles that naturally occurred in our study.

In our population, we find that vertical SPV has roughly the same magnitude as horizontal SPV. Prior studies made at 7T have also observed vertical nystagmus, albeit to a smaller degree (Mian et al., 2016; Otero-Millan et al., 2017). We considered the possibility that the vertical nystagmus we observed was due to a consistent tendency for subjects to have their heads tilted out of alignment with the scanner bore (e.g., due to asymmetric padding in the head coil). Instead, we find no bias in roll angle, and further observe that a roll=0 alignment of the vestibular system with the B_0_ field is still predicted to produce a downward SPV. The relationship between roll and vertical SPV, and the presence of non-zero vertical nystagmus with roll=0 positioning, has been observed previously at 7T when measured in the first 20 seconds of entry into the scanner bore (Mian et al., 2016).

An important caveat to our measurements is that nystagmus (and thus SPV eye movements) can arise spontaneously when people are deprived of visual input. The magnitude and direction of these responses varies with body position, although a downward SPV eye movement is a frequent physiologic response in people placed in a supine position in darkness (Bisdorff et al., 2000). While spontaneous, physiologic nystagmus might contribute to the overall magnitude of vertical SPV eye movements that we have recorded, this mechanism is unable to account for the dependence of vertical SPV upon the roll position of the head with respect to the B_0_ magnetic field.

The cause of MR-induced vertical SPV is unclear. Ward and colleagues postulated that MR-induced Lorentz forces acting upon the superior (aka anterior) semi-circular canals should be (for our polarity of scanner) excitatory on the right, and inhibitory on the left, thus canceling and resulting in zero vertical SPV (Ward et al., 2014). People with unilateral vestibular loss may consequently have a substantial vertical SPV (Ward et al., 2014), and undiagnosed, unilateral vestibular loss in some subjects could contribute to the vertical SPV we measure in our cohort. This account, however, would not explain the particular downward SPV bias that we observe, as it would require a preponderance of right-sided vestibular loss. An alternative explanation is that, due perhaps to the internal configuration of the ampulla, MR-induced endolymph movement is somewhat stronger when directed towards the cupula as opposed to away. Perhaps consistent with this account, a small, *upwards* SPV was consistently observed in measurements made on subjects introduced into a 7T scanner with a B_0_ polarity opposite ours (Otero-Millan et al., 2017). This observation of a reversal in vertical SPV direction with reversal of the polarity of the MRI field also argues against spontaneous, positional nystagmus accounting for these findings across studies.

Our recordings of eye position were made with an infrared camera. The iris appears relatively homogeneous in such images and we did not attempt to extract a measure of torsion (although methods exist that could in principle support such a measurement; Otero-Millan et al., 2015). A torsional SPV movement has been observed previously in studies of MR-induced nystagmus (Otero-Millan et al., 2017; Ward et al., 2019), but we are unable to comment upon the degree or distribution of such effects in our data.

We find that the vestibular signals remain constant across a 20 minute scanning period. Prior studies have shown that there is an initial, rapid adaptation of SPV during the first 200 seconds after entering the magnetic field, followed by a slower decay towards a new, non-zero baseline, with incomplete adaptation to constant acceleration over 90 minutes (Jareonsettasin et al., 2016; Zee et al., 2017). A vertiginous sensation of body rotation is also common when subjects are introduced into the bore of a scanner, but this occurs at higher field strengths than used in this study and typically abates after one or two minutes (Mian et al., 2013).

What effect might vestibular signals and their individual variation have upon resting state BOLD fMRI data collected using HCP protocols? While the neural effect of MR-induced vestibular signals has not been directly measured, imaging studies have examined both the direct effect of vestibular stimulation upon neural activity, and the inferred effect of MR-induced nystagmus upon the correlation of fMRI signals between cortical regions. As reviewed by Zu Eulenburg and colleagues, the cortical site of processing of rotatory vestibular signals is not a settled matter (Zu Eulenburg et al., 2012), but peri-sylvian regions have been consistently identified, including the insula, parietal operculum, and—in a recent positron emission tomography study—Heschel’s gyrus (Boegle et al., 2016). The potential effect of MR-induced vestibular signals upon fMRI data has been examined indirectly. Lorentz forces scale linearly with the strength of the magnetic field. In a series of studies, Boegle and colleagues have examined differences in the structure of resting-state fMRI correlations collected using a 1.5 or 3 Tesla scanner, with the insight that these differences may be attributable to differences in induced vestibular signals (Boegle et al., 2016). They find that the difference in magnetic field strength is related to changes in the correlation between regions of the “default mode network”, and between these regions and the visual cortex. In a more recent study, this group has observe that individual differences in the measured orientation of the inner-ear anatomy of subjects undergoing scanning is related to default mode network correlations with putative vestibular cortical and cerebellar regions (Boegle et al., 2020).

## Conclusion

In summary, we find that persistent slow-phase velocity eye movements are frequently present in subjects studied under standard Human Connectome Project protocols, and that the magnitude of these vestibular signals varies significantly between individuals. We find a consistent vertical SPV that cannot be attributed to a bias in head position, and could be explained by anisotropy in induced endolymph movement. An interesting and practical implication of our measurements is that a slight head tilt may be used to null the population mean vertical SPV. For subjects introduced head-first supine into a 3T Siemens scanner, rolling the head 5-8° towards the right shoulder is predicted to null the mean vertical vestibular signal. This might be accomplished by introducing asymmetric padding into the head coil, or by appropriate design of custom-molded head restraints (Power et al., 2019).

## Author Contributions

Conceptualization: Virginia L. Flanagin and Geoffrey K. Aguirre. Data curation: Giulia Frazzetta, Lauren Cutler, and Saguna Malhotra. Formal analysis: Cammille C. Go, Huseyin O. Taskin, and Seyed-Ahmad Ahmadi. Funding acquisition: Jessica I. Morgan and Geoffrey K. Aguirre. Methodology: Huseyin O. Taskin, Seyed-Ahmad Ahmadi, and Virginia L. Flanagin. Project administration: Jessica I. Morgan and Geoffrey K. Aguirre. Software: Huseyin O. Taskin, Seyed-Ahmad Ahmadi, and Geoffrey K. Aguirre. Supervision: Jessica I. Morgan, Virginia L. Flanagin, and Geoffrey K. Aguirre. Writing - original draft: Cammille C. Go and Geoffrey K. Aguirre. Writing - review & editing: Huseyin O. Taskin, Seyed-Ahmad Ahmadi, Giulia Frazzetta, Lauren Cutler, Saguna Malhotra, Jessica I. Morgan, and Virginia L. Flanagin.

## Study Funding

Supported by National Institutes of Health Grant U01EY025864: Human Connectomes in Low Vision, and P30 EY001583: Core Grant for Vision Research.

## Declaration of Interest

None

## Data/Code Availability Statement

The data described in the article are openly available:

- The T2 images can be found on OpenNeuro: doi:10.18112/openneuro.ds004038.v1.0.0
- Eye tracking videos can be found on YouTube: https://www.youtube.com/channel/UCXojt8KD3bCsUlT_zaWzWWQ

The code used to analyze the data is publicly available in three repositories:

- Analysis of eye tracking videos: https://github.com/gkaguirrelab/eyeTrackTOMEAnalysis/tree/master/code
- Statistical analysis and plotting functions: https://github.com/gkaguirrelab/mriTOMEAnalysis/tree/master/code/innerEarModelAndNystagmus
- Registration of the inner ear template to the T2 images and calculation of semicircular canal: Available at the time of publication

## Notes

https://openneuro.org/datasets/ds004038/versions/1.0.0

https://www.youtube.com/channel/UCXojt8KD3bCsUlT_zaWzWWQ

https://github.com/gkaguirrelab/eyeTrackTOMEAnalysis

https://github.com/gkaguirrelab/mriTOMEAnalysis

### Competing Interest Statement

The authors have declared no competing interest.

